# Modifiable clinical dental impression methods to obtain whole-mouth and detailed dental traits from vertebrates

**DOI:** 10.1101/2023.11.21.567763

**Authors:** Johannes N. Wibisana, Ray A. Sallan, Towa Ota, Pavel Puchenkov, Tai Kubo, Lauren Sallan

**Author notes:** Genomics and Regulatory Systems Unit, Okinawa Institute of Science and Technology, Onna-son, Okinawa, Japan (Johannes N. Wibisana). Center for Data Science, Waseda University, Nishiwaseda, Shinjuku, Tokyo, Japan (Tai Kubo). These authors contributed equally (Johannes N. Wibisana, Ray A. Sallan).

## Abstract

Dental impressions, developed for accurate capture of oral characteristics in human clinical settings, are seldom used in research on non-livestock, non-primate, and especially non-mammalian, vertebrates due to a lack of appropriate tools. Studies of dentitions in most vertebrate species usually require euthanasia and specimen dissection, microCT and other scans with size and resolution tradeoffs, and/or ad-hoc individual impressions or removal of single teeth. These approaches prevent in-vivo studies that factor in growth and other chronological changes and separates teeth from the context of the whole mouth. Here, we describe a non-destructive method for obtaining high resolution dentition-related traits that can be used on both living animals and museum specimens for almost all vertebrates, involving a customizable and printable dental impression tray. This method has repeatedly and accurately capture whole-mouth morphology and detailed features at high resolution in the living non-teleost actinopterygian fish, *Polypterus senegalus,* in a laboratory setting. It can be used for comparative morphology and to observe temporal changes such as the presence of microwear, tooth replacement rates, and occlusal and morphological changes through ontogeny.

**Summary statement:** This study presents a cheap and customizable method using a printable tray to measure oral morphological traits in vertebrates humanely and accurately.

## Introduction

Dentition provides essential information on the ecology and evolution of vertebrate species. In fishes alone, dental traits have been used to investigate wide-ranging topics such as development (Hung et al., 2015), replacement (Carr et al., 2021), dental microwear analysis (Purnell and Darras, 2015), jaw evolution (Ronco et al., 2021), feeding characteristics (Streit et al., 2015), biomechanics and bite force (Deng et al., 2022) and diversification of the oral apparatus (Burress, 2016). However, all these studies used either dead specimen or isolated teeth (Burress, 2016; Streit et al., 2015), and euthanasia was often necessary to obtain samples (Hung et al., 2015; Ronco et al., 2021). Furthermore, dissection of the jaw and/or removal of teeth is typically performed, damaging the original sample. The same constraints limit *in vivo* studies of other vertebrate dentitions and oral traits, apart from those on livestock and a handful of primates for whom clinical dental practices exist (Hoffman et al., 2015; Teaford and Oyen, 1989).

In order to avoid destructive practices in obtaining morphological data, scanning methods (e.g. optical, microCT, and laser microscope) have been increasingly used on full specimens (Akhter and Recker, 2021). However, scanning approaches also have constraints which limit their utility in many vertebrates. Optical scanning, as of 2024, lacks the resolution necessary to see fine details such as wear. MicroCT and Synchrotron X-ray scanning imposes size limitations on specimens and have trade-offs between field of view, processing time, and image resolution, and of course take place under high radiation (Sun et al., 2022). Usage of X-ray scanning therefore requires dead and or dissected samples in most cases and/or considerable preparations and planning at limited facilities for limited *in vivo* work to date (Gradl et al., 2018).

Here, we propose and demonstrate a noninvasive alternative methodology to obtain high resolution dental and oral trait information. Our approach builds on prior work on whole mouth dental morphology in vertebrates, which applied established human clinical dental practices and materials to routinely capture dental morphology in primates (Cuozzo and Sauther, 2006; Teaford et al., 2020, 2017), or used veterinary dental materials to achieve the same in livestock (Hoffman et al., 2015). However, such commercial tools and tested procedures did not exist for most vertebrate species, especially outside mammals where the jaws and dentition may have dimensions significantly different than humans. We designed a method using 3D printed, adjustable dental impression trays which can be used with readily available, inexpensive non-toxic materials to take intraoral records from additional groups of vertebrates.

Prior work has shown that direct application of dental impression material to non-mammalian teeth can accurately capture fine surface details (Purnell and Darras, 2015; Sawaura et al., 2022), suggesting that whole-mouth dental impressioning could provide extensive data on morphological diversity. However, this ad-hoc application in fish and other smaller animals was mostly limited to fixed dead and skeletal specimens or fallen teeth. The use of a species-customized tray allows for repeated use with non-model live animals for tracking chronological changes, and exact capture of whole mouth morphology.

Here, we demonstrate the utility of our modifiable, freely printable tray in taking detailed oral impressions of new species (see below and supplementary figure 1 for download and adjustment details). We developed and tested this methodology on specimens of the non-teleost fish *Polypterus senegalus*. Below, we describe the details of our tray, its proper usage, its potential applications, and the results obtained by this method.

## Materials and methods

### Impression tray design

A dual arch impression tray system (Figure 1A and B) with a handle was chosen as the basis for our design. This will provide both maxillary/upper and mandibular/lower arches in one impression and better utilize sedation time. For obligate air breathers such as tetrapods and some fishes, the custom tray design should account for breathing and steps should be taken to avoid blocking airways with impression material. That can be done by changing external wall height and shape in those areas in addition to careful filling of the upper part of the custom tray with impression material to maintain air flow. We placed a crescent-shaped trench, comparable in size to the width of the teeth (0.5–1 mm for a 15-25 cm *Polypterus* specimen), on both the maxillary and mandibular aspects of the tray. This design allows the dentition to fully submerge in the impression material without touching the tray, ensuring a complete impression of all dental surfaces, including the tips of the teeth (Figure 1A). The tray was designed to enable individual customizations of size and shape using different parameters (Supplementary figure 1).

**Figure 1.**
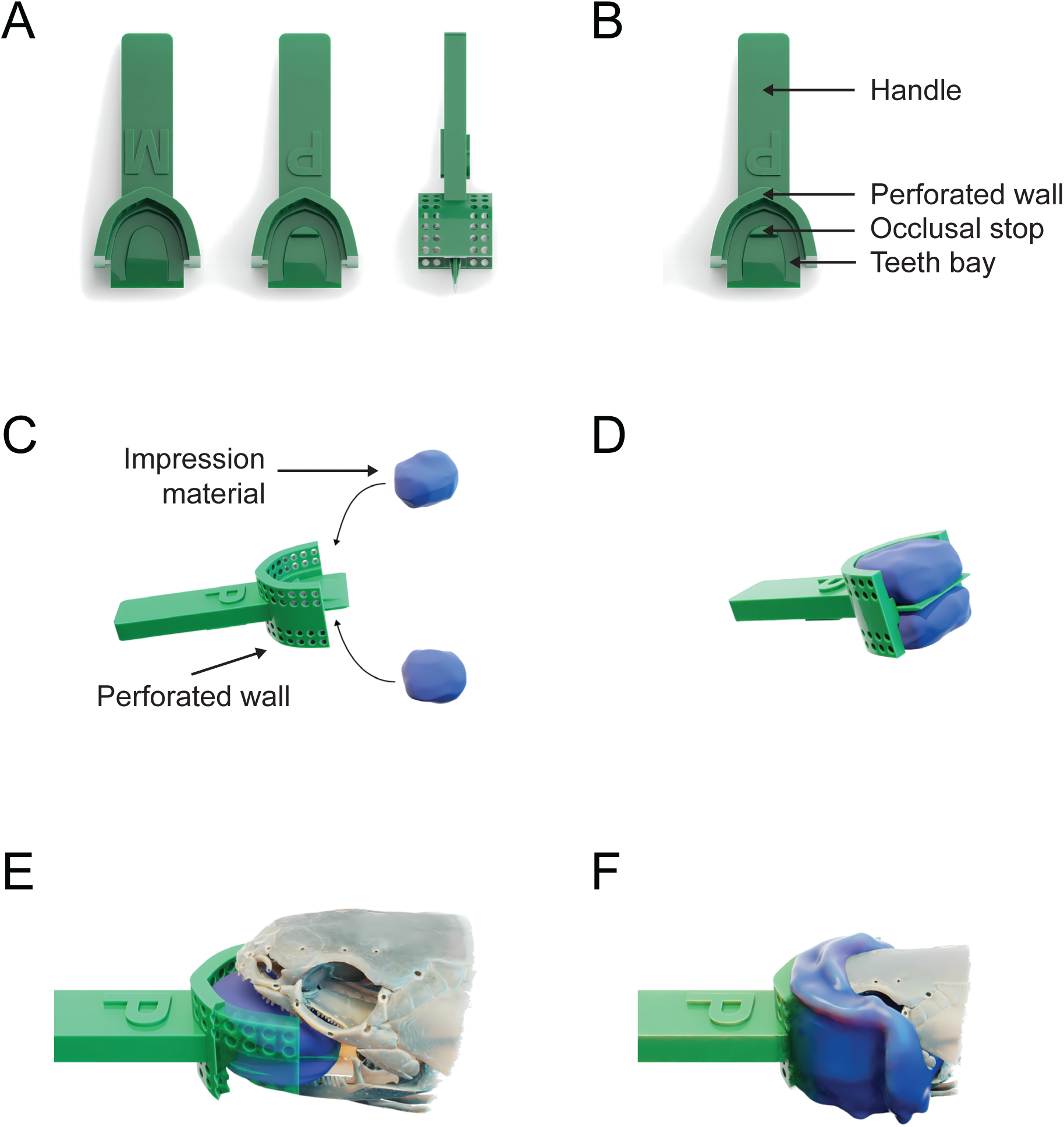
Schematic Representation of the Impression Tray and Usage. Schematic representation of the dental impression tray and impression process, using *Polypterus* as an example. (A) 3D rendered image of the impression tray showing mandibular/lower side, maxillary/upper side, and lateral view. (B) Parts of the impression tray labelled. (C-D) Impression material is applied to both sides of the impression tray. (E-F) *Polypterus* biting on the impression material.

An extraoral perforated retaining wall in front of the trench allows for firm stabilization during impression sequences (Figure 1C-F), as movement by the animal or the operator would otherwise negatively affect accuracy and material distortion. Our wall element extends posteriorly and past to the angle of the mouth in order to affix and stabilize the head. Perforations were included in the design of the retaining wall to aid in attaching the impression material to the tray without the use of adhesives. The height of the tray wall can be customized to match the external border and size of the jaw (Figure 1A, right). The inner aspects of the tray include a raised portion to act as a bite stop (or an occlusal stop) preventing the tray from touching the teeth (Figure 1A, middle). Incorporating these stops, which can be placed based on oral morphology, ensures a uniform thickness of impression material (Terry et al., 2010), increases accuracy, and minimizes distortion (Ortensi and Strocchi, 2000). In our *Polypterus* test animals, the centrally located parasphenoid bone on the oral roof and the “tongue” (basihyal) in the mandibular floor were designated as hard/soft tissue stops. We inserted a centralized raised stop behind the trench on maxillary section of the tray and a smaller raised central lingual floor on the mandibular side in our test trays, both of which can be removed before printing (Supplementary figure 1).

The final version of our adjustable, 3D-printable impression trays was designed using Autodesk Fusion360 and adjustable parameters were added using Blender 4.0. Inclusion of sliders and checkboxes in Blender to adjust the size, relative dimensions, and intraoral and extraoral features (stops, holders, etc.) enables application to a wide range of mouth shapes and allows the tray to be modified without technical skills in 3D modeling. The model is freely available and archived on Zenodo (doi: 10.5281/zenodo.12524788). The steps to modify the tray are outlined in supplementary figure 1.

### Fish husbandry

*Polypterus senegalus* Cuvier specimens of 20-25cm size of random sex were acquired from local pet shops. 100 L water tubs with Eheim Classic 2215 external filters (Eheim, Germany) are used to house up to 2 *Polypterus senegalus* Cuvier specimens of 20-25cm size. The water temperature is controlled to be 25°C. Feeding is performed 3 times a week using 2 pellets per fish of Hikari Crest Freak Bottoms (Kyorin, Japan), however any other food for bottom dwelling carnivorous fish should be admissible. 10-20% water change is performed every week, removing detritus and uneaten food. Water quality, including pH, nitrate and nitrite contents are monitored after every feeding, changing 10-20% of water when nitrate or nitrite levels show higher than around 25ppm and 1ppm respectively.

### Fish sedation

1:2 ratio by mass of tricaine mesylate (MS-222) (Sigma-Aldrich, USA) and NaHCO_3_ (Nacalai Tesque, Japan) as buffer were used for fish sedation, following previous reported sedation of *Polypterus* (Whitlow et al., 2022). Each fish was transferred to a smaller tank of 2 L of water with dissolved buffered MS-222 (Figure 2B). Optimization of fish sedation was performed by gradually changing the MS-222 concentration in increments of 50 mg L^-1^, starting from 100 mg L^-1^ with 5 minutes of exposure, prior to transferring each fish outside the water, and measuring the time before it starts moving. We performed these trials using 3 different fishes per session and recording the shortest time for any fish to recover. We performed this trial once per day, to let the fish recover upon sedation. For approximately 25 cm *Polypterus senegalus* Cuvier, 5 minutes of exposure to 300 mg L-1 of MS-222 was enough to keep the fish sedated for at least 5 minutes. We used 1.5 mg L-1 of MS-222 to keep the fish sedated for 30 minutes, enough to obtain 5 impressions.

**Figure 2.**
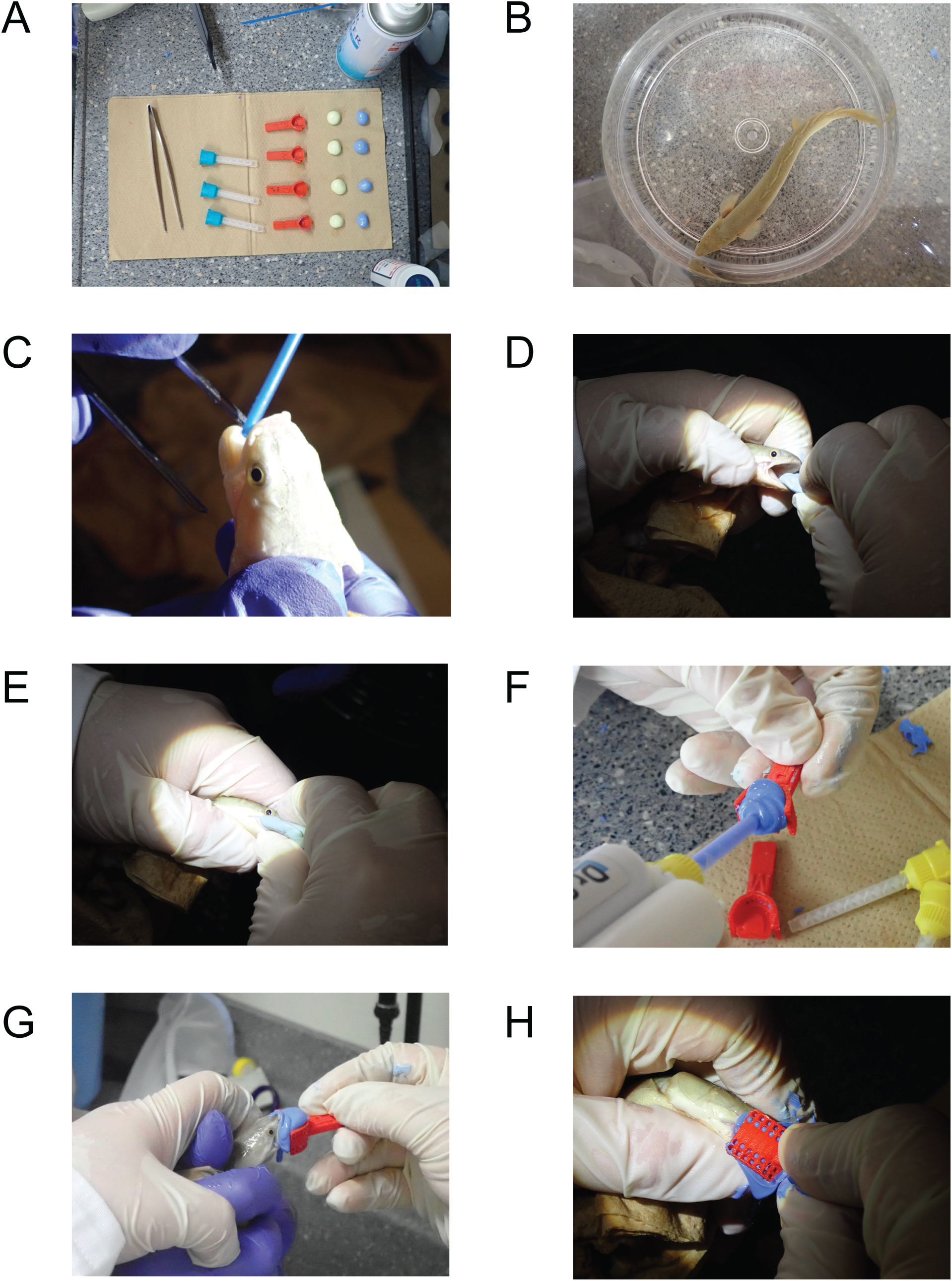
Example Impression Sequence Using *Polypterus*. Molding sequence of a 20 cm *Polypterus* with a ∼1.4 cm head width, 1.7 cm head length and 1.5 cm gape. (A) Preparation of materials before starting the procedure. (B) Individual in buffered MS-222 bath. (C) Removal of residual moisture using compressed air duster. (D-E) Removal of impurities from teeth surface using putty. (F) Application of PVS material on the surface of the impression tray. (G-H) Molding of the teeth by firmly pushing the jaw onto the impression tray. The fingers of the operator are used to firmly push the jaws together from the top and below, the other hand of the operator is used to hold the impression tray in place.

### Intraoral impression process

The following protocol was developed through testing with live and preserved *Polypterus* species and can be adapted for other species using similar impression materials. The tray design can be modified based on the oral morphology of the animal. We recommend taking an initial bite with rigid and less expensive material, such as PVS putty, during the planning process to assess general oral size and features, which will aid in tray design. The protocol for sedation for all animals should be developed in consultation with a veterinarian, the animal care and use committee, and/or established published guidelines.

### Scanning of dental impressions

Obtained dental impressions were scanned using a confocal laser microscope (VK-X210, Keyence, Japan) with a violet laser (408nm) and an ultra-long range 100x lens. Molds were cut along the line connecting mesio-distal section of each tooth using a scalpel and dental loupes (Supplementary figure 2), so that both buccal/labial and lingual/palatal surfaces of teeth can be observed in the impression. The tips of teeth were scanned as this area contact most intensively with food and likely have more dental microwear on the basis of work with other non-mammalian tetrapods (Winkler et al., 2019). The three-dimensional surface models and two-dimensional images were generated using the photo-simulation function of the laser microscope.

### Adjusting and printing impression trays

We first adjust our tray models using Blender to the specifications for species and individuals. We then 3D print trays using an Ultimaker S5 (Ultimaker, Netherlands) with Ultimaker PLA material (Ultimaker, Netherlands), but any printer can be used with PLA material from any source. Ultimaker Cura is used to slice the STL files generated from Blender, using the default preset “extra fine” with 0.06 mm layer height. We find that the tray prints best with the support overhang angle at 80° and brim build plate adhesion type, printing the structure upright with the handle on top and the mouthpiece at the bottom. Less Z axis resolution can be used with larger impression trays. The size of the tray is mostly limited by the build volume of the 3D printer, in our case 33×24×30cm. However, for even larger animals, trays can be printed in several different parts by splitting the 3D model and manually reassembling it post-print.

### First impression to remove debris

After preparation (Figure 2A) and confirming that the animal is fully sedated (Figure 2B), tweezers were carefully used to gently pry the mouth open to expose the oral cavity. Gentle compressed air is applied for 3–5 seconds to dry the dentition and disperse any residual moisture which can be aquatic or salivary in nature (Figure 2C). This step is crucial as moisture has a negative effect on the accuracy of PVS impressions (Walker et al., 2005). The first impression is taken using Vinyl Polysiloxane Putty Type (ExaFine, GC, Japan). Following the manufacturer’s recommended mixing process, the putty can either be applied directly to the dentition, or the animal can be gently made to bite into it 2–3 times while the putty is relatively soft (Figures 2D-E). The putty should be removed about 1 minute prior to complete hardening to minimize the risk of enamel damage.

### Final impression sequence

To prepare for the final impression using a more accurate impression material, an Automix tip is affixed to a Polyvinyl siloxane (PVS) cartridge (Dr. Silicon regular, BSA Sakurai, Japan) to ensure proper mixing of impression components. This material has been shown to cure quickly and has high accuracy (Sawaura et al., 2022), suitable for a wide range of applications including dental microwear study, and is routinely available from dental suppliers (Supplementary table 1). A portion of the impression material is exuded from the tip prior to filling the impression tray to ensure that only properly mixed material is injected. To avoid air pockets, the impression dispenser Automix tip remains submerged inside the impression material as it is ejected into the tray (Figure 2F).

The loaded impression tray is placed into the mouth of the animal, where both jaws are pressed together to simulate a bite (Figure 2G). Animal and impression tray are held together securely yet gently to avoid harm until the impression material is completely set (Figure 2H). The curing time was set to 5 minutes to allow the material to harden or fully cure. This duration is optimal for medium-body silicone impression material at room temperature (Sawaura et al., 2022). Our own testing of the curing duration for the Dr. Silicon impression system on *Polypterus* yielded similar results. Depending on the manufacturer, using lighter or thicker viscosity impression materials may require different curing times. It is important to remember that these dental impression systems are designed for the warm-blooded human intraoral environment and that is reflected in the manufacturers’ directions. They will harden more quickly inside a mammalian mouth due to the temperature differences. In contrast, when used extraorally or in a cooler intraoral environment, such as that of fish, the material hardens at a slower rate. In addition to material testing, it is advisable to consult the manufacturer’s specifications prior to experimenting on animals. The live animal is then returned to its original confines and monitored until it regains motor functions. This two-impression process takes approximately 10 minutes.

### Geometric Morphometric Analysis

We imaged the dental impressions using Hirox digital microscope HRX-01 (Hirox, Japan) with a high-range motorized zoom lens 20x-2500x. As there are no standardized oral soft or hard tissue landmarks for *Polypterus*, fourteen set landmarks and ten semi-landmarked curves were chosen around the parasphenoid and ectopterygoid bones, representing common and reproducible points on different impressions. Two-dimensional morphometric data was collected from the digitized dental impressions in R version 4.3.1 using the StereoMorph package v1.6.7 (Olsen and Westneat, 2015). Procrustes distances and PCA analyses were subsequently performed using the package borealis v2022.10.27 (https://github.com/aphanotus/borealis). Disparity is calculated as the Procrustes variance using the diagonal elements from the group covariance matrix divided by the number of observations and was derived using the function “morphol.disparity()” from the package geomorph v4.0.7 (Baken et al., 2021). The codes and raw data used for this analysis are available and archived on Zenodo (doi: 10.5281/zenodo.12524788).

### Ethics statement

This research was approved by the ethics committee of the Okinawa Institute of Science and Technology under the protocol number ACUP-2023-006-2.

## Results

### Trait resolution (Based on *Polypterus)*

Using this method, we were able to produce impressions that accurately captured dentition and intraoral morphology of very small live specimens of *Polypterus* (Figure 3A-B). We scanned the impressions taken from different individuals (see Materials and Methods for details of scanning and image production) and effectively captured dental structures from around 1 micron to 1 mm across the entire jaw (Figure 3C-D, Supplementary figures 3A-C), a higher resolution than a previous study using direct manual measurements (Mihalitsis and Bellwood, 2019). From 38 impression attempts on 12 individuals, there was no incident of specimen mortality or injury due to the impression process.

**Figure 3.**
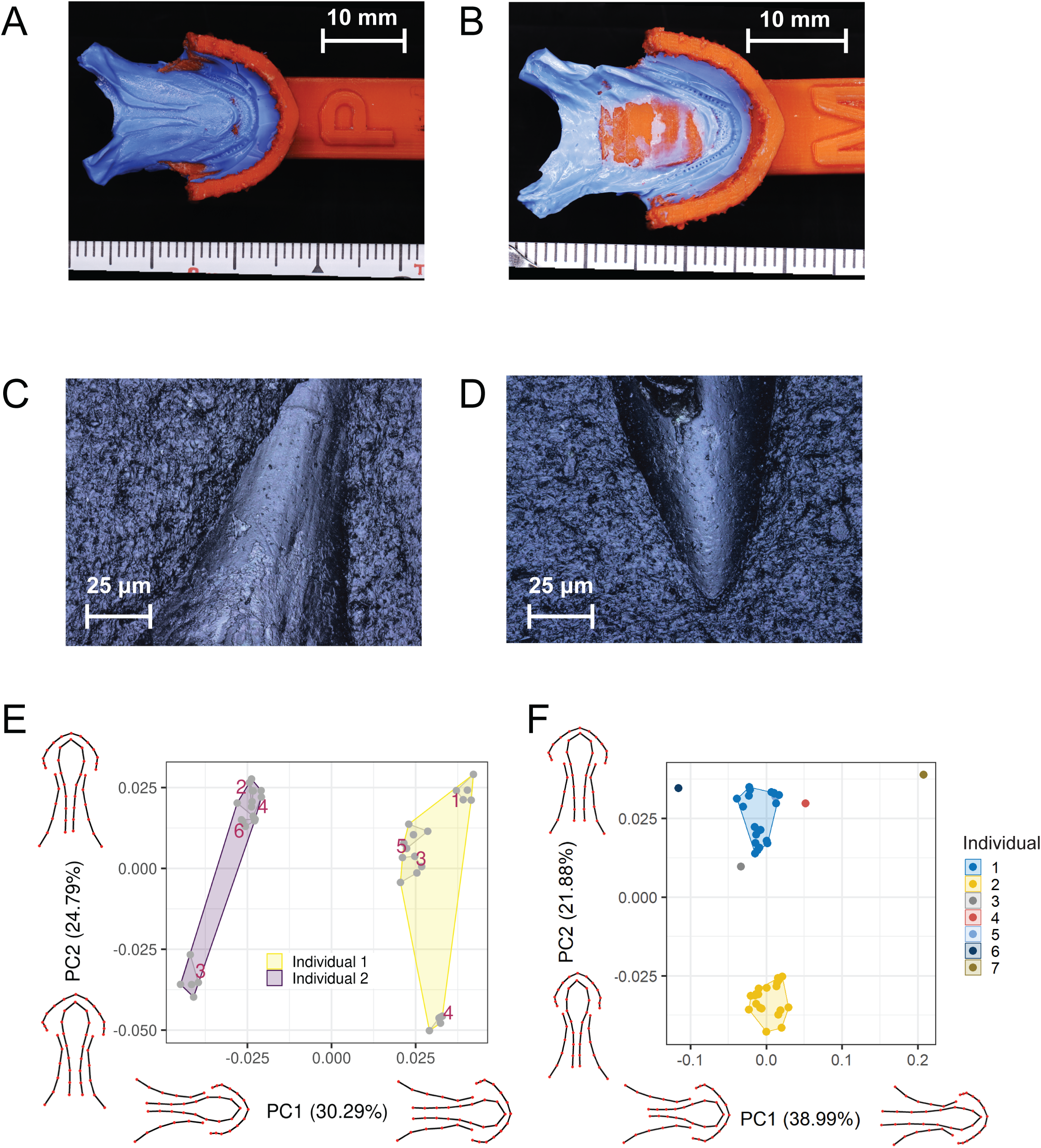
Example impressions from *Polypterus*. PVS mold obtained upon the completion of impressions. (A-B) Undetached impression from (A) maxillary side and (B) mandibular side. (C-D) Lingual scan of impression obtained from the maxillary side for (C) posterior tooth and (D) anterior tooth. (E) PCA plot of geometrics morphometrics analysis using landmarks from 4 maxillary/upper impressions of each of 2 individuals. 5 replicate landmark sets were digitized from each impression (represented by red convex hulls). Trial numbers are indicated over each hull group. (F) PCA plot of geometrics morphometrics analysis of all available whole mouth impressions (N=13) from 7 individuals used in method development. Some impressions were omitted when they were deemed to be deformed due to contact between animal and tray (eliminating putty) and could not be landmarked, or when they were cut to obtain details on tooth impressions featured in Fig. 3A.

Because of the small size of our specimens (15-25 cm total length with a bite length of 1.5 cm) and their teeth (<1 mm in width), it was difficult to conduct Dental Microwear Texture Analysis (DMTA), as true dental microwear may have been similar in size to the limits of putty and laser scanner resolution and therefore hard to distinguish. In addition, all our test animals were fed only soft pellets, limiting the amount of dental microwear produced by grit or hard material. However, we identified enamel damage on some teeth at the resolution of ∼10 microns by quantifying the depths of the ridges on the surface of the teeth (Figure 3C-D, supplementary figures 3A-B). It is likely that our method can capture more widespread microwear from larger teeth for DMTA, as this has been captured from direct application of putty to molar-like teeth from other species (Purnell and Darras, 2015). Prior DMTA studies have all been on larger teeth, such as those from mammals and lizards (Teaford et al., 2020; Winkler et al., 2022).

### Reproducibility

As this is a qualitative method, we tried to assess the reproducibility and accuracy of obtaining shape information through geometric morphometric analysis of maxillary/upper impressions (Figure 3E-F, Supplementary figures 4-5). This allowed quantification of impression method error, landmark assignment operator error, and individual variation, based on generating 5 sets of landmarks for the parasphenoid and associated soft tissues in each of 4 maxillary impressions on each of 2 different, very small (approximately 25 cm length) individuals in one sedation session (40 total landmark sets) (Supplementary figure 4, 5A-C). We performed principal component analysis (PCA) (Figure 3E, Supplementary figures 5B-C), and found that PC1 was associated with differences between individuals, while PC2 was associated with some impression method error (the difference between a tight clustering of most impressions and an outlier). Outliers were not related to order of impression and did not include any initial impressions. Using the same parasphenoid landmarks, we also conducted a geometric morphometrics analysis of all available whole maxillary impressions from 7 individuals taken over multiple sessions, spread over the multi-month course of method development and data collection for another project (Figure 3F, Supplementary figures 5D-E). Our PCA results for all individuals further demonstrated that variation between different individuals or the same individual’s growth over time was much greater than any method error or operator error generated in a single session. This suggests that this method can quantify the gross oral morphology of individual specimens reproducibly, and capture variation over time within individuals and across populations.

We calculated disparity values for landmark sets from duplicate images of single impressions, multiple sequential impressions of one individual, and from all 7 study individuals (Supplementary figure 5F). Levels of variance between duplicate images of one impression were comparable, representing similar levels of operator error between observations. Disparity was highest for all impressions from individuals (reflective of individual variation) and lower for multiple impressions from one individual even including outliers on PC2, reflective of method error and not the sequence in which impressions were taken. This indicates the method can typically accurately detect and represent distinctions between individual variations with high fidelity and a low rate of error in shape.

## Discussion

As above, this method can be used to non-destructively and cheaply obtain whole mouth oral traits at high resolution from both live and dead animals for which commercially available dental trays are unsuitable, such as fish, reptiles, and mammals outside primates and domesticated families. It can be used for any morphological studies involving oral traits, occlusion, and dentition in place of existing methods that are destructive (e.g. dissection), expensive (e.g. microCT), as well as methods with less control (e.g. direct application of putty), or reduced resolution (e.g, optical scanning). This method has achieved resolution of structures down to at least 1 micron, such as enameloid damage from feeding and differences in enameloid ultrastructure (e.g. the difference between acrodine and other enameloid in actinopterygian fishes).

One of the most important benefits of this method is that it allows a simplified process of obtaining time series data of detailed morphological changes in dentition and oral-pharyngeal cavity in animals such as fish. This is useful for studies of dental microwear analysis (Hoffman et al., 2015) as well as growth, replacement, and development (Carr et al., 2021; Streelman et al., 2003; Trapani et al., 2005). It can be used to record detailed intraoral soft-tissue anatomy of living animals including features that previously could only be evaluated on dead specimens (Luczkovich et al., 1995). While the silicone impression method has been used previously, particularly on primates, the modification described in this paper provides the practicality of creating a cost-effective and customizable impression tray. This new approach allows for adaptation to a wide range of animals, offering significant potential for broader applications in oral morphological studies.

### Potential applications

Some possible applications of this tray-mediated impression method include the production of physical and 3D digital models from live or rare specimens which cannot be prepared for high resolution scans by microCT or other means. Impressions can be used for geometric morphometric analyses of dentitions in a range of species. Occlusion patterns preserved in impressions can provide an alternative approach to study bite articulation and jaw biomechanics (Wilga and Ferry, 2015). Soft tissue and high resolution dental models created from scans of impressions can also be of use with the physical replication of skeletal movements and feeding mechanics for (XROMM) studies of individuals, which typically depend on whole body CT scans at low resolution (Camp and Brainerd, 2015). Although tooth replacement has been extensively studied from a histological and developmental perspective there is very little data on tooth growth and replacement on living non-mammal individuals (Brink et al., 2021; Carr et al., 2021; Ellis et al., 2015), especially with time series from a single individual. The recurring creation of whole-dentition tooth records via dental impression provides an opportunity to not only study replacement patterns but also examine external factors that can affect it. We are positive that researchers focused on the ecology and evolution of vertebrates will find many additional uses for the data captured by our method.

### Material sourcing and cost considerations

Costs are reduced by using existing, consumer-grade, and readily available 3D printers and material. In principle, most rigid 3D printing materials can be used, here we opted to use poly-lactic acid as it is one of the most available and cheap materials. Dental impression compounds can also be selected based on the purpose. For fish, the use of MS-222 anesthetic can be costly, thus clove oil can be used as a substitute due to its potency and cost (Javahery et al., 2012). The same anesthesia mix can also be reused for multiple aquatic individuals, as it is mediated by water. The breakdown of the materials and costs for our tests are available in supplementary table 1.

## Method limitations and considerations

### Sedation of live animals

Changes to size or species might require modification of the sedation method to accommodate time required for material polymerization. Air breathing *Polypterus* has an advantage in surviving outside of water. When applying this method to other aquatic animals, physiology is crucial in determining survivability and thus should be thoroughly investigated. Due to the hydrophilic nature of PVS, the body can be submerged under water and only the jaw exposed to air during the impression process

Maintenance of sedation might be required when more time is needed to take impressions, while optimization of anesthesia concentration should be performed depending on species and size. If longer time is required, maintenance of sedation by running oxygenated water with anesthetics through the gills might be required.

### Use with air breathing species

For obligate air breathers such as tetrapods and some fishes, the custom tray design should account for breathing and steps should be taken to avoid blocking airways with impression material. In addition to carefully and conservatively filling the impression tray, another technique is to position or tilt the head and body downward during the impression process. This helps the material flow away from the throat area. The tray can also be modified to accommodate airway breathing by adjusting the height and shape of the external walls in the nasal areas. Additionally, a distal lip can be added to the inner edge of the impression tray to prevent material overflow, but it should not hinder occlusion.

### Mitigating the enamel pellicle and biofilm

In-vivo dental microwear analysis is complicated by pellicle (protein film) formation on enamel, which can affect impression accuracy and obscure microwear details (Hoffman et al., 2015). Operational time constraints necessitated that we have a quick and safe process to remove biofilm prior to impression. Using diluted bleach can have negative effects on PVS material (Hamalian et al., 2011) and teeth (Sim et al., 2001), while brushing can deform enamel surface (Wiegand et al., 2007). Therefore, we opted for a dual impression technique to remove biofilm.(Hamalian et al., 2011; Sim et al., 2001; Wiegand et al., 2007)

### Tooth and animal size

PVS dental impression materials are mainly used to precisely replicate human teeth with high reproducibility. In the case of smaller oral morphology, molding success becomes technique-dependent, making reproducibility challenging. Therefore, it is recommended to obtain impressions from multiple specimens, and repeatedly on a single specimen to determine the best material and confirm accuracy in different species. Further, morphometric data or external photos or optical scans of specimens can be used to help modify the custom trays for other species and individual animals, reducing the need for iterative printing and testing.

### Pharyngeal and extraoral teeth

It is possible to produce pharyngeal teeth impressions with customization of the tray to extend intraorally, and we have included options for this within our model. However, precautions must be taken to avoid injury or ingestion of the impression material. Due to the lack of deep pharyngeal teeth, this could not be tested with *Polypterus.* However, internal gill morphology deep within the pharynx was captured alongside the marginal dentition (Figure 3A-B)

### Interpretation of soft tissue results in preserved specimens

While this method can be easily applied to dead and preserved specimens, users are advised to use caution in the interpretation of soft-tissue dependent whole mouth traits such as occlusion patterns. It is likely that post-mortem and preservational changes can affect the movement of muscles, allowing the jaws to move into positions that are not possible in life.

### Use with species with kinetic and complex bites

Many vertebrates, such as teleost fishes, have protrusible jaws or other kinetic dental and jaw elements. Impressioning these species requires further validation and possibly introduces additional limitations. One concern is the accuracy of occlusal patterns captured by our method, as operator manipulation of the jaw in a sedated animal might produce unnatural or uneven bite patterns. However, at least in teleosts, we expect that, since peak force is applied at the end of the bite (Westneat, 2003), and there is little lateral jaw movement in most fishes, occlusion patterns would be constrained and therefore similar for *in vivo* molds of the same animal. However, as above, it is possible muscle changes in preserved specimens could change movement patterns.

The degree of occlusal variation and bite flexibility in species with kinetic jaws mechanics can and should be tested by comparing multiple impressions from one individual, in the same way we took multiple sequential impressions of sedated *Polypterus* to determine overall accuracy (Figure 3). Based on operator knowledge of the jaw kinetics of the target species, the tray can also be adjusted in Blender to add sloping external walls to guide the bite. Until species-specific validation is performed, we suggest that the focus should be on morphological features of the upper and lower jaw in species with kinetic bites, rather than occlusal patterns. This is also because most such species do not use manipulative or biting feeding modes, but rather suction or ram (Westneat, 2003).

## Acknowledgments

We thank the OIST Animal Resource Section and Diala Joy Edde for the support provided in fish husbandry and impressions. We are grateful for the 3D printing support provided by the Machine Engineering section and the 3D visualization support provided by the Scientific Computing and Data Analysis Section of the Core Facilities at Okinawa Institute of Science and Technology Graduate University. We thank Chloe Nash for advice on morphospace analysis.

## Data availability

Parameterized blender file and STL files for the impression trays used in this project are available on https://anonymous.4open.science/r/impressiontray-24FE/README.md and is archived on Zenodo (doi:10.5281/zenodo.12524788).

## Author contributions

J.N.W., R.S., T.K., and L.S. conceived the project. J.N.W. and R.S. conceived the method and performed overall direction and planning. R.S. created the initial tray design. J.N.W., T.K., and R.S., performed molding trials. J.N.W. designed and fabricated 3D models of dental trays. P.P parameterized and provided 3D visualization for the impression tray, molding and cutting sequence. T.K. performed microscope scanning of obtained molds. T.O. performed morphometric analyses. J.N.W., T.K., R.S. and L.S. wrote the manuscript. All authors contributed critically to the drafts and gave final approval for publication.

---

**Supplementary Figure 1.**
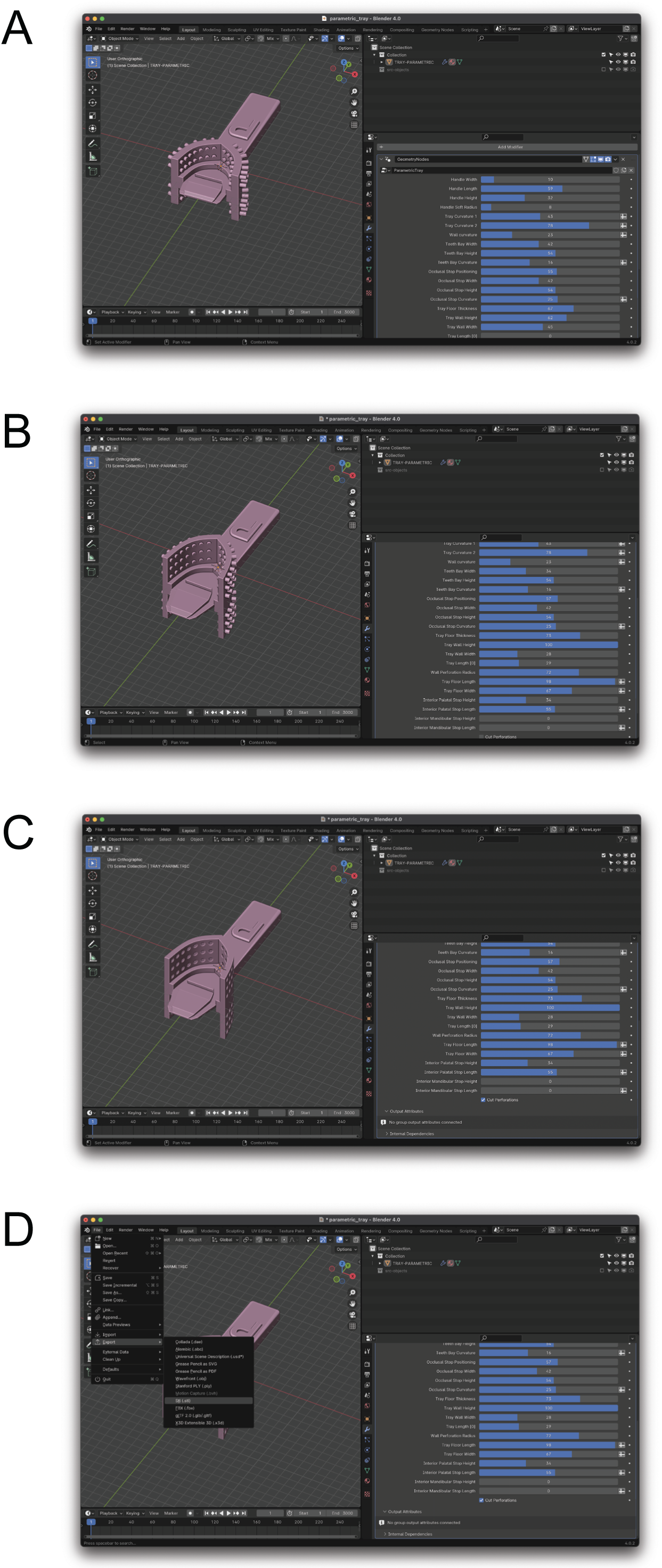
Tray customization. Customization of the impression tray using Blender. (A) The supplementary (or downloadable) file parametric_tray.blend can be loaded into Blender. (B) Relative parameters can be adjusted depending on the target species by adjusting the values or the slider on the pane on the right side of the window. (C) Perforations need to be cut after adjustments and before exporting to STL as it is a computationally intensive process. (D) STL can be exported by selecting export > Stl (.stl).

**Supplementary Figure 2.**
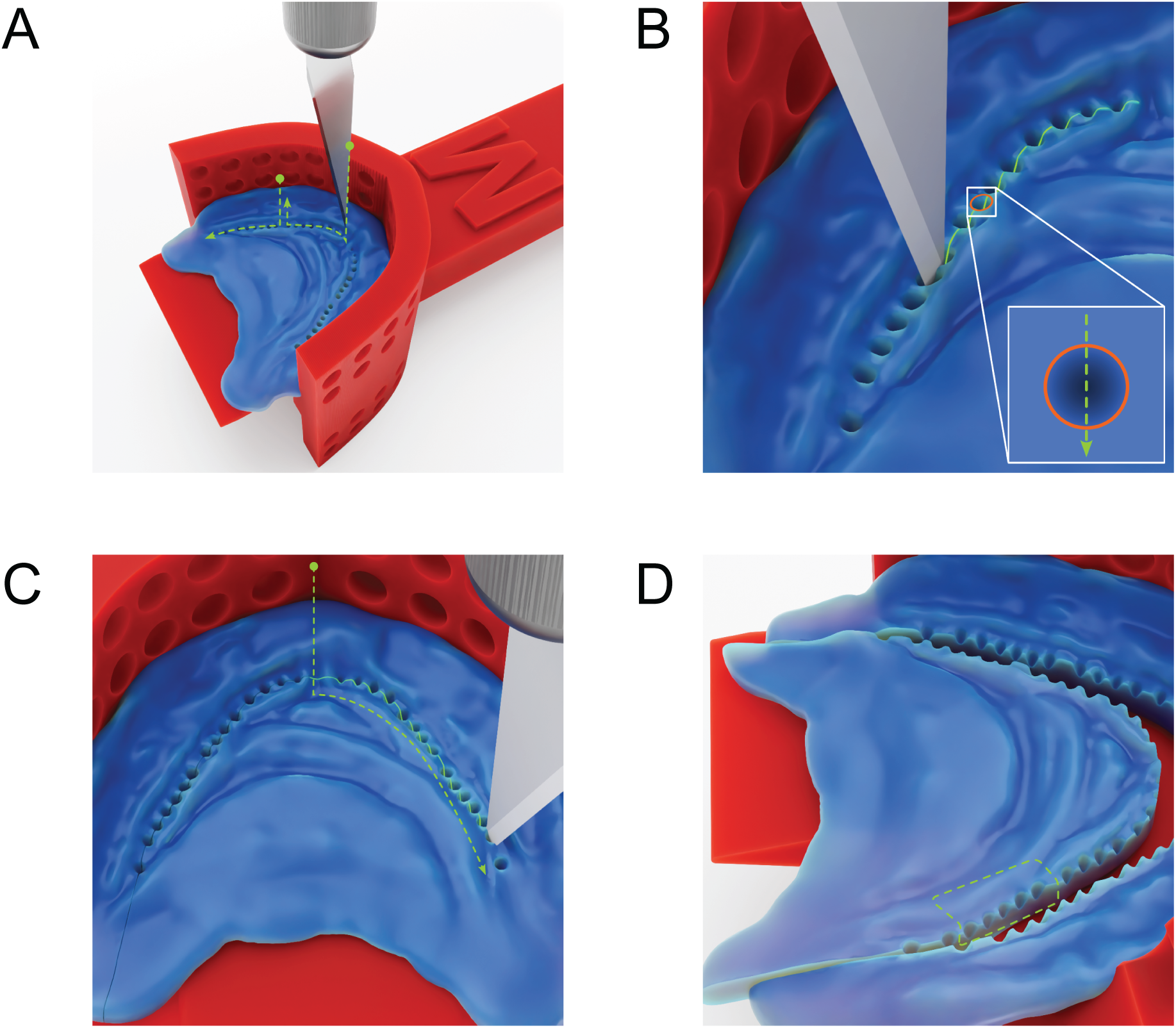
Processing of impression before scanning. 3D rendering of steps taken to obtain a scannable impression. (A) Scalpel was used to cut along the dentition holes. (B) Close-up image of (A). (C) Cutting was performed along the dentition holes until the ends. (D) The result of the impression being cut in half.

**Supplementary Figure 2.**
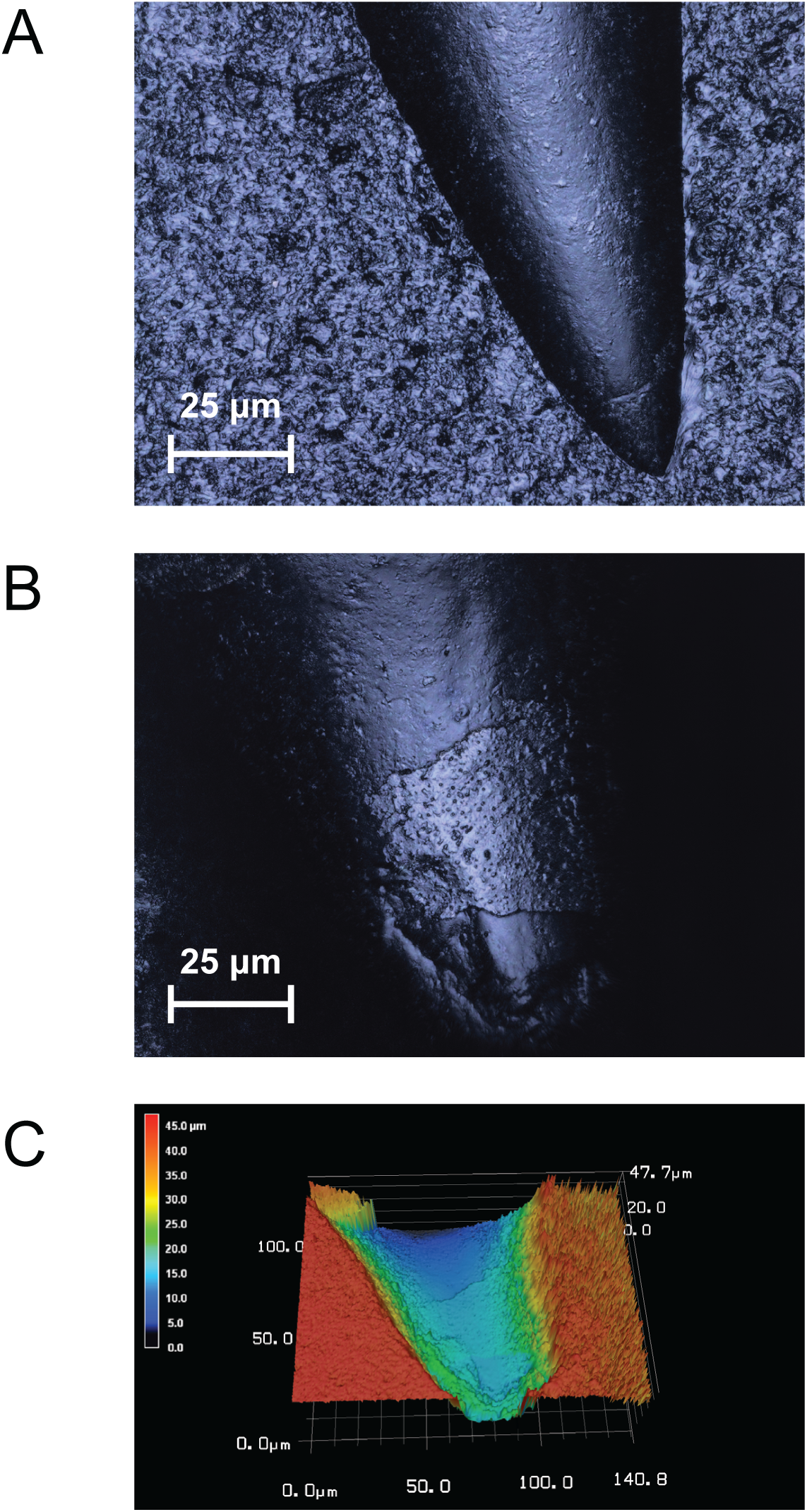
3D scanning of tooth impression. (A) the mandibular side for posterior tooth. (B-C) Lingual scan of the mandibular side for a middle tooth, which enamel was flaked off by brushing, (B) photo-simulation and (C) topology showing the elevation at the area where enamel was flaked off. The size of photo-simulations (A-B) is 100 micrometers (Y axis) times 140.5 micrometers (X axis).

**Supplementary Figure 4.**
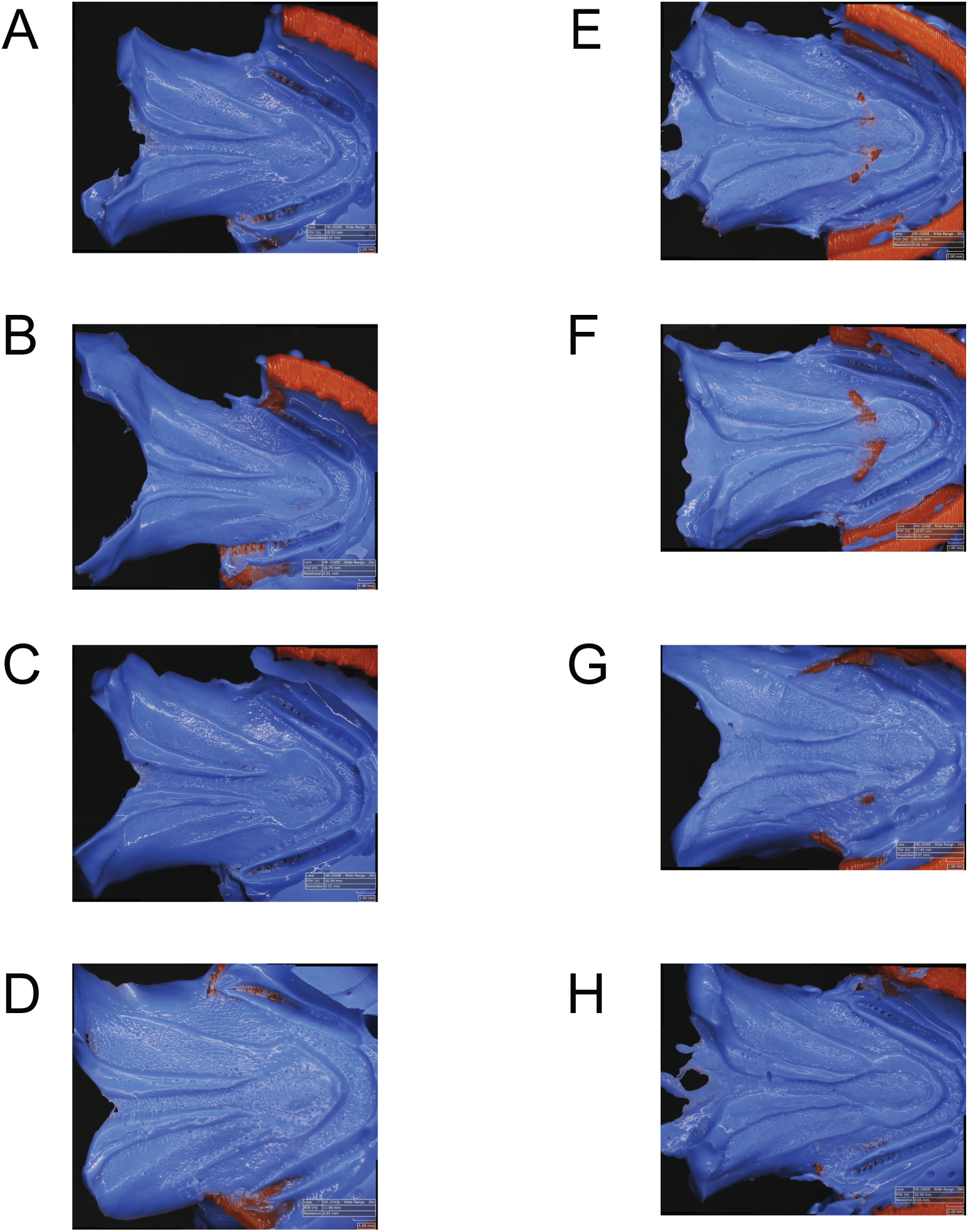
Impression taken from two individuals for morphometrics analysis. Impressions obtained from 2 different individuals in single sedation sessions (∼20 minutes total). (A-D) Impressions taken from individual 1 representing trials 1, 3, 4, 5 respectively. (E-H) Impressions taken from individual 2 representing trials 2, 3, 4, 6 respectively.

**Supplementary Figure 5.**
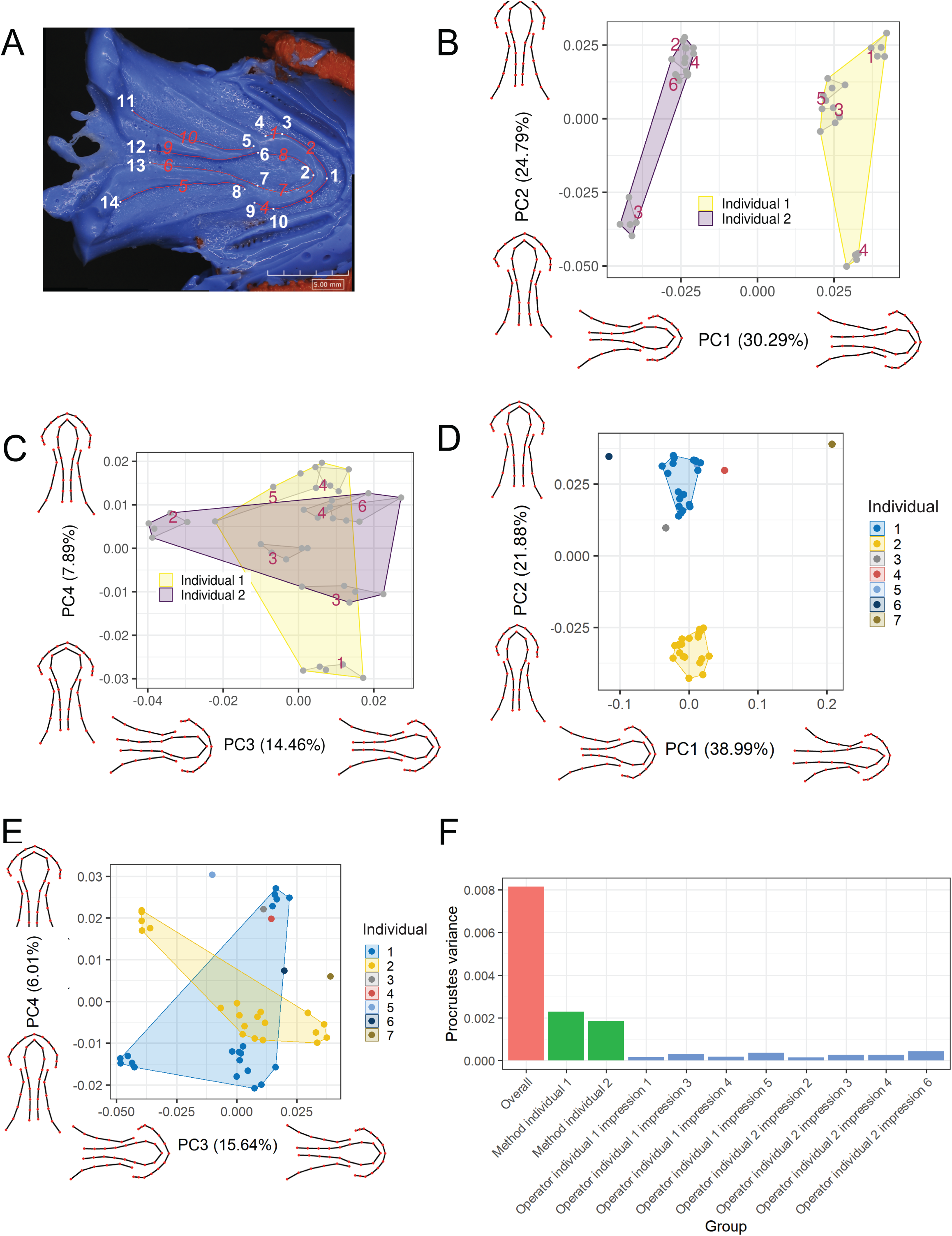
Geometric morphometrics analysis. Geometric morphometrics analyses. (A) Landmarks used to perform morphometrics analysis. 42 landmarks were placed in total; white numbers are fixed landmarks, and red numbers are semi-landmarked curves. (B-C) PCA plot (N=40) showing (B) PC1 and PC2 and (C) PC3 and PC4 from the morphometrics analysis of 2 individuals. Each impression is represented by the red convex hull. Trial numbers are indicated over each hull group. (D-E) PCA plot (N=65) showing (D) PC1 and PC2 and (E) PC3 and PC4 from the morphometrics analysis of 7 different individuals. (F) Disparity calculations for image duplicates and impression duplicates. “Overall” disparity shows procrustes variance across impressions taken from individuals 1-7. “Method” disparity shows procrustes variance between all impressions taken from specimen 1 and specimen 2 respectively. “Operator” disparity shows the difference between all duplicated images from each impression used in the principal components and represents operator error.

**Supplementary Table 1.** Impression material cost. The approximate cost of impression material for the experiments performed in (A) this study and (B) alternative cost for impression materials with a more cost-effective option.

| Type | Item | Manufacturer | Approximate cost in USD | Use per experiment | Approximate cost per experiment in USD | Note |
| --- | --- | --- | --- | --- | --- | --- |
| Impression material | Dr. Silicone regular type (medium body) 50ml cartridges | BSA sakurai | 10 per cartridge | ~5ml per animal | 1 | Tray size 2 cm by 2 cm deep and included filling palatal and mandibular sides. |
| Impression material | Exifone putty 556 ml (combined base and catalyst) | GC Japan | 60 | 2.8 ml per animal | 0.3 | A quarter of a mix of 1 scoop of base and 1 scoop of catalyst. |
| Anesthesia | Ethyl m-Aminobenzoate Methanesulfonate (MS-222) 25g | Nacalai Tesque | 108 | 0.6g in 2L of water | 2.6 | Anesthetic mixed water can be used for multiple fish cohabiting the same tank. |
| 3D printing | UltiMaker NFC PLA Filament 2.85mm (750g) | Ulti Maker | 50 | 1g | 0.1 | Autoclavable trays can be reused on different animals. |

**Supplementary Table 2.** Alternative impression material cost. The approximate alternative cost for impression materials with a more cost-effective option. Alternative materials can also be used when performing experiments on larger fishes that require higher anesthesia dose.

| Type | Item | Manufacturer | Approximate cost in USD | Use per experiment | Approximate cost per experiment in USD | Note |
| --- | --- | --- | --- | --- | --- | --- |
| Anesthesia | Clove oil C8392-100ML | Sigma Aldrich | 55 | Approximately less than 0.2ml | 0.1 | We assume that ethanol that is used to dissolve the clove oil is already present in most labs |
| 3D printing | Generic PLA 1kg | various | 20-30 | 1g | less than 0.1 |  |
| 3D printing | Generic ABS 1kg | various | 20-30 | 1g | less than 0.1 |  |
| 3D printing | Generic PETG 1kg | various | 20-30 | 1g | less than 0.1 |  |
| 3D printing service | 3D printing service | various | 1-2 | - | 1-2 |  |

## References

Akhter MP, Recker RR. 2021. High resolution imaging in bone tissue research-review. Bone 143:115620. doi:10.1016/j.bone.2020.115620

Baken EK, Collyer ML, Kaliontzopoulou A, Adams DC. 2021. geomorph v4.0 and gmShiny: Enhanced analytics and a new graphical interface for a comprehensive morphometric experience. Methods Ecol Evol 12:2355–2363. doi:10.1111/2041-210X.13723

Burress ED. 2016. Ecological diversification associated with the pharyngeal jaw diversity of Neotropical cichlid fishes. J Anim Ecol 85:302–313. doi:10.1111/1365-2656.12457

Camp AL, Brainerd EL. 2015. Reevaluating Musculoskeletal Linkages in Suction-Feeding Fishes with X-Ray Reconstruction of Moving Morphology (XROMM). Integr Comp Biol 55:36–47. doi:10.1093/icb/icv034

Carr EM, Summers AP, Cohen KE. 2021. The moment of tooth: rate, fate and pattern of Pacific lingcod dentition revealed by pulse-chase. Proc Biol Sci 288:20211436. doi:10.1098/rspb.2021.1436

Cuozzo FP, Sauther ML. 2006. Severe wear and tooth loss in wild ring-tailed lemurs (Lemur catta): A function of feeding ecology, dental structure, and individual life history. J Hum Evol 51:490–505. doi:10.1016/j.jhevol.2006.07.001

Deng Z, Loh H-C, Jia Z, Stifler CA, Masic A, Gilbert PUPA, Shahar R, Li L. 2022. Black Drum Fish Teeth: Built for Crushing Mollusk Shells. Acta Biomater 137:147–161. doi:10.1016/j.actbio.2021.10.023

Gradl R, Dierolf M, Günther B, Hehn L, Möller W, Kutschke D, Yang L, Donnelley M, Murrie R, Erl A, Stoeger T, Gleich B, Achterhold K, Schmid O, Pfeiffer F, Morgan KS. 2018. In vivo Dynamic Phase-Contrast X-ray Imaging using a Compact Light Source. Sci Rep 8:6788. doi:10.1038/s41598-018-24763-8

Hamalian TA, Nasr E, Chidiac JJ. 2011. Impression Materials in Fixed Prosthodontics: Influence of Choice on Clinical Procedure: Impression Materials: A Review. J Prosthodont 20:153–160. doi:10.1111/j.1532-849X.2010.00673.x

Hoffman JM, Fraser D, Clementz MT. 2015. Controlled feeding trials with ungulates: a new application of *in vivo* dental molding to assess the abrasive factors of microwear. J Exp Biol. doi:10.1242/jeb.118406

Hung NM, Ryan TM, Stauffer JR, Madsen H. 2015. Does hardness of food affect the development of pharyngeal teeth of the black carp, Mylopharyngodon piceus (Pisces: Cyprinidae)? Biol Control 80:156–159. doi:10.1016/j.biocontrol.2014.10.001

Javahery S, Nekoubin H, Moradlu AH. 2012. Effect of anaesthesia with clove oil in fish (review). Fish Physiol Biochem 38:1545–1552. doi:10.1007/s10695-012-9682-5

Luczkovich JJ, Norton SR, Gilmore RG. 1995. The influence of oral anatomy on prey selection during the ontogeny of two percoid fishes, Lagodon rhomboides and Centropomus undecimalis. Environ Biol Fishes 44:79–95. doi:10.1007/BF00005908

Mihalitsis M, Bellwood D. 2019. Functional implications of dentition-based morphotypes in piscivorous fishes. R Soc Open Sci 6:190040. doi:10.1098/rsos.190040

Olsen AM, Westneat MW. 2015. StereoMorph: an R package for the collection of 3D landmarks and curves using a stereo camera set-up. Methods Ecol Evol 6:351–356. doi:10.1111/2041-210X.12326

Ortensi L, Strocchi ML. 2000. Modified custom tray. J Prosthet Dent 84:237–240. doi:10.1067/mpr.2000.108453

Purnell MA, Darras LPG. 2015. 3D tooth microwear texture analysis in fishes as a test of dietary hypotheses of durophagy. Surf Topogr Metrol Prop 4:014006. doi:10.1088/2051-672X/4/1/014006

Ronco F, Matschiner M, Böhne A, Boila A, Büscher HH, El Taher A, Indermaur A, Malinsky M, Ricci V, Kahmen A, Jentoft S, Salzburger W. 2021. Drivers and dynamics of a massive adaptive radiation in cichlid fishes. Nature 589:76–81. doi:10.1038/s41586-020-2930-4

Sawaura R, Kimura Y, Kubo MO. 2022. Accuracy of dental microwear impressions by physical properties of silicone materials. Front Ecol Evol 10. doi:10.3389/fevo.2022.975283

Sim TPC, Knowles JC, Ng Y-L, Shelton J, Gulabivala K. 2001. Effect of sodium hypochlorite on mechanical properties of dentine and tooth surface strain: Effect of sodium hypochlorite on dentine. Int Endod J 34:120–132. doi:10.1046/j.1365-2591.2001.00357.x

Streelman JT, Webb JF, Albertson RC, Kocher TD. 2003. The cusp of evolution and development: a model of cichlid tooth shape diversity. Evol Dev 5:600–608. doi:10.1046/j.1525-142x.2003.03065.x

Streit RP, Hoey AS, Bellwood DR. 2015. Feeding characteristics reveal functional distinctions among browsing herbivorous fishes on coral reefs. Coral Reefs 34:1037–1047. doi:10.1007/s00338-015-1322-y

Sun R, Wang Y, Zhang J, Deng T, Yi Q, Yu B, Huang M, Li G, Jiang X. 2022. Synchrotron radiation X-ray imaging with large field of view and high resolution using micro-scanning method. J Synchrotron Radiat 29:1241–1250. doi:10.1107/S1600577522007652

Teaford MF, Oyen OJ. 1989. Live primates and dental replication: New problems and new techniques. Am J Phys Anthropol 80:73–81. doi:10.1002/ajpa.1330800109

Teaford MF, Ungar PS, Taylor AB, Ross CF, Vinyard CJ. 2020. The dental microwear of hard-object feeding in laboratory *Sapajus apella* and its implications for dental microwear formation. Am J Phys Anthropol 171:439–455. doi:10.1002/ajpa.24000

Teaford MF, Ungar PS, Taylor AB, Ross CF, Vinyard CJ. 2017. In vivo rates of dental microwear formation in laboratory primates fed different food items. Biosurface Biotribology 3:166–173. doi:10.1016/j.bsbt.2017.11.005

Terry DA, Tric O, Blatz M, Burgess JO. 2010. The custom impression tray: fabrication and utilization. Dent Today 29:132, 134–5.

Trapani J, Yamamoto Y, Stock DW. 2005. Ontogenetic transition from unicuspid to multicuspid oral dentition in a teleost fish: Astyanax mexicanus, the Mexican tetra (Ostariophysi: Characidae). Zool J Linn Soc 145:523–538. doi:10.1111/j.1096-3642.2005.00193.x

Walker MP, Petrie CS, Haj-Ali R, Spencer P, Dumas C, Williams K. 2005. Moisture Effect on Polyether and Polyvinylsiloxane Dimensional Accuracy and Detail Reproduction. J Prosthodont 14:158–163. doi:10.1111/j.1532-849X.2005.04024.x

Westneat MW. 2003. A biomechanical model for analysis of muscle force, power output and lower jaw motion in fishes. J Theor Biol 223:269–281. doi:10.1016/S0022-5193(03)00058-4

Whitlow KR, Ross CF, Gidmark NJ, Laurence-Chasen JD, Westneat MW. 2022. Suction feeding biomechanics of *Polypterus bichir*J: investigating linkage mechanisms and the contributions of cranial kinesis to oral cavity volume change. J Exp Biol 225. doi:10.1242/jeb.243283

Wiegand A, Wegehaupt F, Werner C, Attin T. 2007. Susceptibility of acid-softened enamel to mechanical wear--ultrasonication versus toothbrushing abrasion. Caries Res 41:56–60. doi:10.1159/000096106

Wilga CAD, Ferry LA. 2015. Functional Anatomy and Biomechanics of Feeding in ElasmobranchsFish Physiology. Elsevier. pp. 153–187. doi:10.1016/B978-0-12-801289-5.00004-3

Winkler DE, Iijima M, Blob RW, Kubo T, Kubo MO. 2022. Controlled feeding experiments with juvenile alligators reveal microscopic dental wear texture patterns associated with hard-object feeding. Front Ecol Evol 10:957725. doi:10.3389/fevo.2022.957725

Winkler DE, Schulz-Kornas E, Kaiser TM, Tütken T. 2019. Dental microwear texture reflects dietary tendencies in extant Lepidosauria despite their limited use of oral food processing. Proc R Soc B Biol Sci 286:20190544. doi:10.1098/rspb.2019.0544

